# Genotypic and phenotypic characterization of *Streptococcus mutans* isolated from dental caries

**DOI:** 10.1101/2020.04.20.050781

**Authors:** Md. Shahadat Hossain, Sadab Alam, Yead Morshed Nibir, Tahrima Arman Tusty, Sayyeed Mahmud Bulbul, Shahidul Islam, Mohammad Shahnoor Hossain

## Abstract

*Streptococcus mutans*, considered as principal causative agent of dental caries, maintains a biofilm lifestyle in the dental plaque. The oral cavity harbors numerous *S. mutans* strains, which displayed remarkable genotypic and phenotypic diversity. This study evaluated the genotypic and phenotypic diversity of 209 *S. mutans* strains isolated from 336 patients with dental caries and compared with the universal reference strain UA159. Our study has revealed a high degree of genotypic and phenotypic variability among the clinical strains. We observed significant differences in colony morphology, generation time, biofilm formation, bacteriocin and acid production while growing in culture medium. All the clinical isolates were able to lower pH while growing in THY broth. In consistent with phenotypic variations, we also observed tremendous level of genotypic variation by AP-PCR and gene specific PCR. AP-PCR analysis suggested that most of the patients with dental caries have distinct type of *S. mutans* strains. Genes related to various two component systems were highly conserved among the strains, however, bacteriocin encoding genes such as *nlmAB, nlmC* were absent in half of the clinical isolates. In sum, our study highlights the genotypic and phenotypic diversity of *S. mutans* clinical isolates and indicates the presence of diverse mechanism to initiate and establish the biofilm lifestyle which leads to tooth decay.

## Introduction

The initiation and successful development of dental caries is caused by multiple bacterial and host factors, such as the composition and biochemical activity of the biofilm organisms, dietary habit, genetic constitution and behavior of the host, tooth architecture and exposure to fluoride (1–4). The mutans streptococci, specifically *S. mutans*, are considered to be the primary causative agents of dental caries, commonly known as tooth decay (5, 6). In addition, several recent studies have reported the associations of certain sub-groups of *S. mutans* with cardiovascular disease (7–11). The ability to form biofilm on tooth surface, production of organic acid from various carbohydrates (acidogenicity), ability to survive at low pH (acidurance), outstanding ability to outcompete other bacteria by the production of bacteriocin and their adaptability to rapidly changing environment can be attributed as the major virulence factors (12–18). Development of natural competence, which is coordinately regulated with the bacteriocin production, is another vital attribute that provides genetic diversity to *S. mutans* for niche adaptation and colonization (19–21).

Strains belong to the species, *S. mutans*, are generally classified into four (c, e, f, and k) serological groups based on the composition of cell-surface rhamose-glucose polysaccharides (22, 23). Strains belonged to serotype c are the most abundant in the oral cavity (70–80%), followed by serotype e (20%) and serotype f or k (2–5%) (1, 24). However, serotype k is more prevalent in heart valves and atheromatous plaques (12%) than in the oral cavity (11). Previously, several attempts have been made to correlate the caries incidence with certain genotypes of *S. mutans*, however, no co-relation was observed among multiple studies (21, 25–28). Furthermore, no relationship was found between the caries status of an individual and the distribution of 41 putative virulence genes or genetic elements in 33 *S. mutans* isolates (29). Nonetheless, several genes have been identified as the virulence attributes and a connection with virulence has been indicated by experiments based on gene inactivation followed by *in vitro* assays (30–33) or by virulence testing in animal models (34–36).

Restriction fragments length polymorphism (RFLP) based fingerprinting (37, 38), multilocus sequence typing (MLST) (39, 40), comparative genome hybridization (41) and the comparison of whole genome sequencing (42, 43) have revealed the prominent intraspecies genetic variability of *S. mutans*. Additionally, several studies have demonstrated the genetic variability of *S. mutans* in individual genes (44, 45). *S. mutans* strains also display phenotypic variability in accordance with the variation in their genetic repertoire (24, 28, 46, 47). This is especially important for *S. mutans*, which is naturally competent bacterium and therefore has the potential for rapid genome diversification through horizontal gene transfer (48). In a previous study, Palmer et al. observed a high degree of phenotypic variability among 15 of the completed draft genomes of 57 geographically and genetically diverse isolates of *S. mutans* (1). Nevertheless, further studies are necessary to get more insights into the genotypic and phenotypic variation among clinically relevant *S. mutans* isolates.

The aim of the present study was to investigate the genotypic and phenotypic heterogeneity of *S. mutans* isolates from 336 patients with dental caries from Bangladesh. We found that *S. mutans* strains isolated from dental caries have high level of genotypic and phenotypic heterogeneity.

## Materials and Methods

### Study population

Samples were collected from 336 of different age and sex groups with dental caries in this study. The study protocol for human subjects was approved by the Institutional Review Board of the Faculty of Biological Sciences, University of Dhaka (Ref. No. 82/Biol.Scs). An informed written consent was taken from each participant. General physiological information of the patients was collected by interview. All the patients did not have any chronic diseases. Patients who had taken antibiotic therapy for the last two weeks were excluded from the study.

### Bacterial culture and growth

Oral samples were collected from patients with dental caries using sterile toothpick and suspended in 1 ml phosphate buffer saline (PBS) buffer. 100 μl of the sample was spread on the Mitis Salivarius agar supplemented with 0.5 IU/mL bacitracin (Sigma, USA) and incubated at microaerophilic condition at 37°C for 48 hours. All strains were stored in 40% glycerol at −80°C and freshly streaked on THY agar before each experiment. *Streptococcus* strains were routinely grown in Todd-Hewitt medium (HiMedia, India) supplemented with 0.2% yeast extract (THY) at 37°C. For biofilm assay, strains were grown in a THY medium supplemented with 1% sucrose. For the monitoring of growth, overnight cultures were diluted into fresh medium (1:20), grown to late exponential phase (OD_600_ = 0.5) and absorbance was taken at 630 nm at various time interval.

### Identification of *S. mutans* strains

Colony morphology on the mitis salivarius-bacitracin agar medium (MSB) was primarily used for the selection of *S. mutans* (49) and confirmed by PCR with species specific primers (Table-4) (50). Three colonies of *S. mutans* per individual were selected from Todd-Hewitt agar plate and preserved at −80°C for later genotypic and phenotypic characterization. However, a single isolate from each patient was investigated in this study. Species-specific PCR (Smu.479) was performed by colony PCR. Briefly, a single colony was picked from the THY agar plate and the cells were suspended directly into PCR mixture in a microcentrifuge tube. The PCR assay included 30 cycles of denaturing at 95°C for 30 seconds, annealing at 50°C for 45 seconds and extension at 72°C for 1 min. The amplicon, generated from PCR reaction, was run in 1.5% agarose gel containing ethidium bromide and checked for the appropriate bands under UV transilluminator.

### Typing of clinical isolates by AP-PCR

The genetic diversity of *S. mutans* isolates was analyzed by arbitrarily primed PCR (AP-PCR) by using the primers set OPA 02 (5’-TGCCGAGCTG-3’) and OPA 13 (5’-CAGCACCCAC-3’) as described previously (21). The colony PCR was performed using 2X PCR master mix under the conditions of 35 cycles of denaturation at 94°C for 30 seconds, annealing at 32°C for 1.5 minutes, extension at 72 °C for 2 minutes, with initial denaturation at 95°C for 5 minutes, and a final extension at 72°C for 10 minutes. The amplicons generated by AP-PCR were then separated by 1.5 % agarose gel electrophoresis. The molecular size of each bands were calculated and a dendrogram was generated using the UPGMA cluster analysis and analyzed by using the Dice coefficient (>95%) in accordance with Mitchell et al. (51).

### PCR amplification of virulence genes

The detection of *nlmA, nlmB, nlmC*, smu.925, *comC,comD comE, gtfB, gbpA, vicK, ciaH, cnm, cbp, atp*, and Smu.1906 was performed by colony PCR using primers specific to gene based on the UA159 genome sequence (Table-4). In addition to the strains being tested, purified genomic DNA from *S. mutans* UA159 was used as a positive control and distilled water was used as a negative control in each PCR. The PCR products were analyzed by electrophoresis in a 1.5% agarose gel. *gyrA* gene was used as an internal control.

### Biofilm assay

For biofilm assay, overnight grown bacteria in THY broth were diluted 1:20 and inoculated into fresh THY medium supplemented with 1% sucrose into wells of polystyrene flat-bottom 24-well microtiter plate and incubated for 48-hr at 37°C in microaerophilic condition. After incubation, the culture medium was decanted, and the wells were washed thrice with distilled water and stained with 1% crystal violet for ten minutes. The plates were further washed with distilled water twice to remove the unabsorbed dye. The cells were then resuspended into 1-ml 95% ethyl alcohol and absorbance was taken at 550 nm. Each experiment was performed in triplicates.

### Investigation of the acidurity of *S. mutans*

To investigate the acidurity of *S. mutans* strains, overnight grown bacterial cultures were diluted to 1:20 into THY broth as control or THY broth acidified with HCl (pH 5.0) in 96 well microtiter plate and incubated at 37°C and the growth was monitored for 24-hour at 630 nm using a microplate reader (Micro Read 1000, ELISA plate analyzer, Global Diagnostics, Belgium).

### Acidogenesis of *S. mutans* clinical isolates

While growing on sugar, *S. mutans* produces acid which, in turn, reduces the pH that causes tooth decay (17). In order to investigate the acid production capacity of the clinical isolates, overnight grown *S. mutans* culture was inoculated in 10 ml of THY broth (pH 8.32) and the pH was determined at different time intervals (0-hr, 24-hr, 48-hr and 72-hr) using a pH meter. Each experiment was performed in duplicate.

### Deferred antagonism bacteriocin assay

To investigate the bacteriocin production by the clinical isolates, isolated colonies were stabbed into THY agar plates with a toothpick and grown overnight (~18-hr) at 37°C under microaerophilic conditions. Indicator strains were grown to mid exponential growth phase in THY broth and 0.4 ml of the indicator culture (*S. pyogenes* and *Lactococcus lactis.)* was mixed with 10-ml of soft agar and overlaid on agar plates that were stabbed with the tester strains. Overlaid plates were then incubated overnight under same conditions and the diameter of the zones of inhibition around the mutacin-producing strains was measured. The isolates were recorded as bacteriocin producer against the indicator bacteria if the zone of diameter was 5-mm or greater.

## Results

### Dental health analysis of diabetes

The DMFT (decay-missing-filled-Teeth) index is widely used to assess the epidemiology of dental caries status. The mean DMFT values of this study was 5.2 ±2.8 and 4.4±2.5 for male and female respectively (n=336) with the mean age of 42.4 and 38.6 for male and female respectively. Among the 336 patients we recruited, 61.6 % was male and 38.4% female. Among the study subjects, 18% were diabetic. Physiological characteristics of the patients with colonization of *S. mutans* are presented in Table 1.

**Table 1:** Physiological description of the study population.

| Characteristics |  | Male | Female |
| --- | --- | --- | --- |
| Number |  | 207 (61.6%) | 129 (38.4%) |
| Age group | 5 to 15 | 23 | 8 |
|  | 15 to 30 | 41 | 17 |
|  | >30 | 143 | 104 |
| Mean Age |  | 42.4 | 38.6 |
| DMFT index | | $5.2 \pm 2.8$ | $4.4 \pm 2.5$ |
| Type II diabetes |  | 45 | 14 |

### Isolation and identification of *S. mutans* clinical strains

In our study, we used mitis-salivarius-bacitracin (MSB) agar medium to isolate *S. mutans* strains due to high selectivity of this medium for *streptococcus*. Based on different colony morphology on MSB medium, colonies were selected for further study. Morphological and cultural characteristics of the isolates ranged from unduly shaped, round sized, blue colonies with granular frosted glass appearance to round, blue, and rough and shiny colonies (data not shown). There were also round or spherical form, raised or convex elevation and black or blue color ranging from a pinpoint to pinhead size with a rough surface, flat, light blue or dark colonies on the MSB agar plate. Pinpoint colony with granular, frosted glass appearance was primarily selected as *S. mutans*. All the strains displayed positive gram staining reaction and catalase negative (data not shown). However, they exhibited either alpha or gamma hemolysis pattern on blood agar (data not shown). Colony PCR with species-specific primers further confirmed the isolates as *S. mutans* since the expected product size of SMU.479 was found in agarose gel after electrophoresis (data not shown). The prevalence of *S. mutans* was 82.14% in patients suffering from dental caries.76% of the preliminary identified colonies were finally confirmed as *S. mutans*. The summary of the results for various characteristics of selected 30 strains has been presented in Table 2.

**Table 2.**
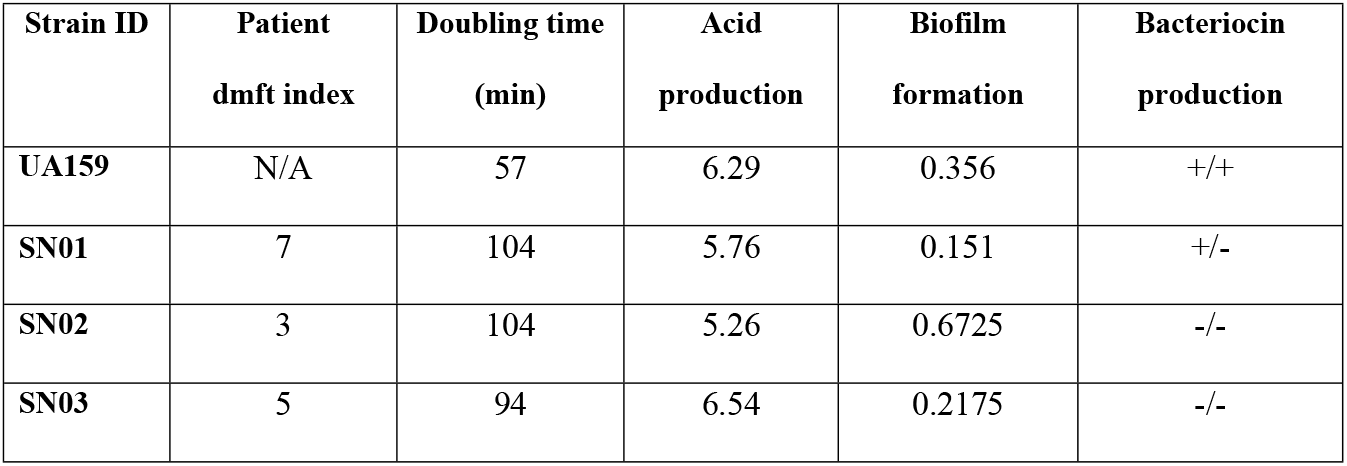
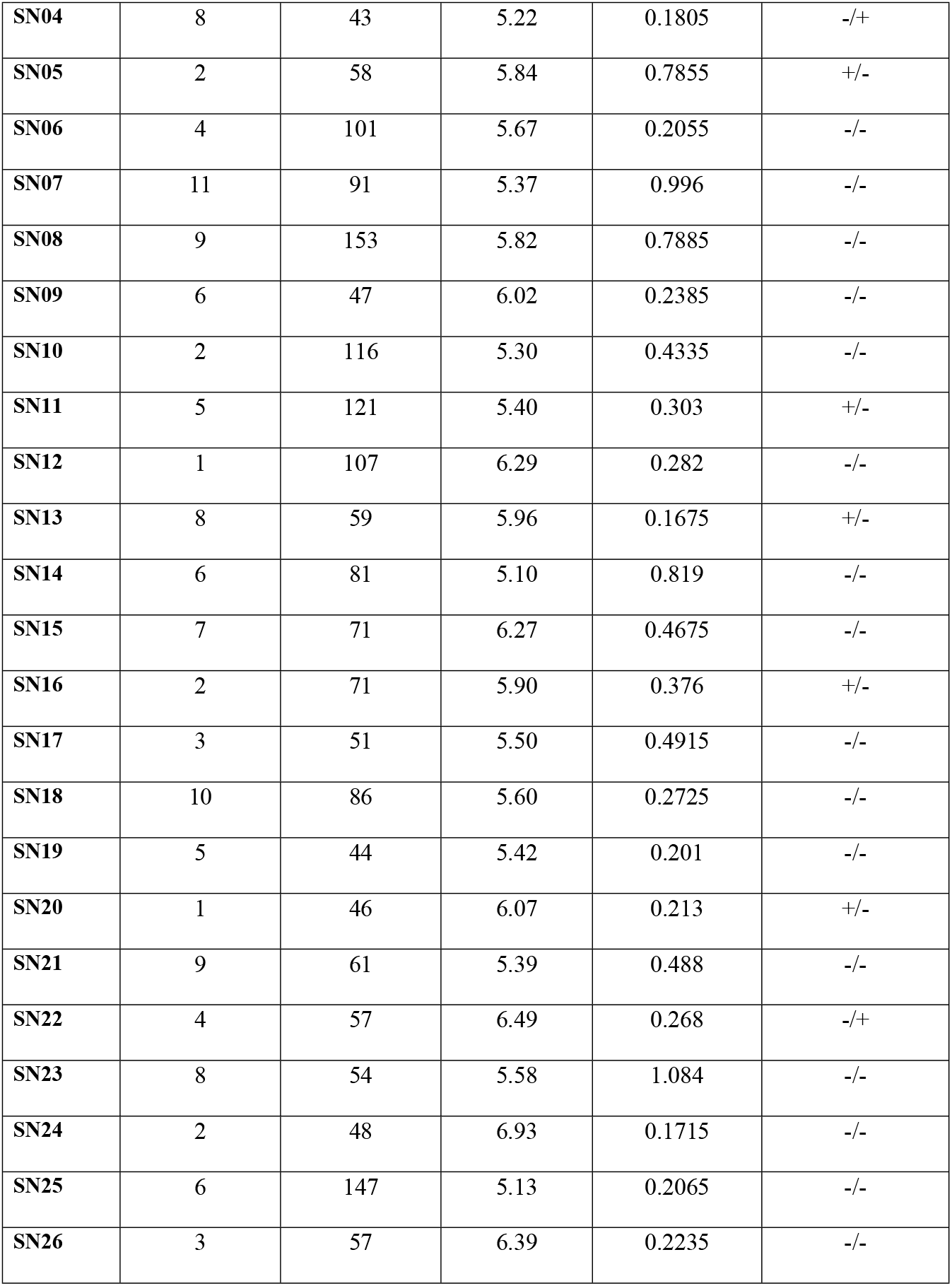
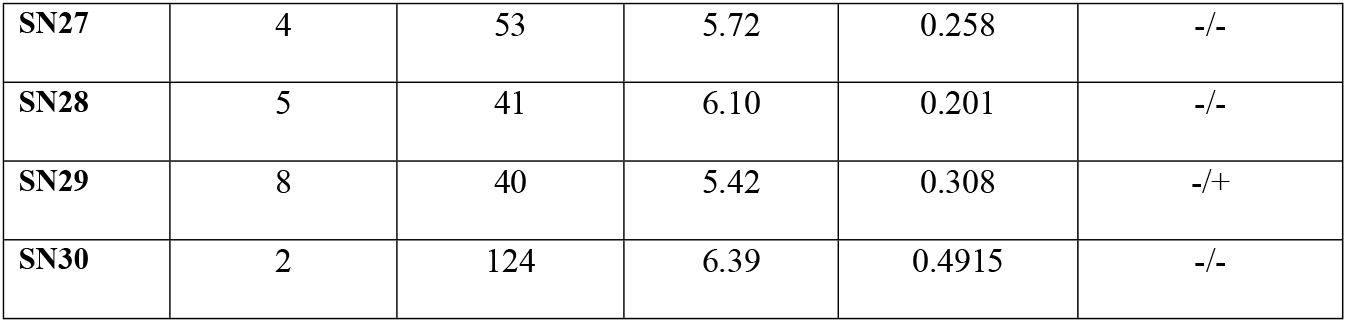
Summary of the results for various phenotypic characteristics of selected strains.

### Genotypic diversity of *S. mutans* isolated

A total of 209 isolates of *S. mutans* were selected for further genotyping assay from the patients with dental caries. Fig 1a and Fig 1b demonstrates the AP-PCR patterns carried out with OPA-02 and OPA-13 primers using some representative isolates, where we observed different spectrum of amplicons for each isolate, which indicates the high level of genetic polymorphism among the isolated strains. The results of AP-PCR analysis of the selected 40 isolates revealed that 26 different genotypes were present among the strains. After analysis of dendrogram of the selected strains, 10 different clusters were observed in our study. No significant correlation of *S. mutans* was observed between genotypic diversity of *S. mutans* in respect to age or gender.

**Fig 1.**
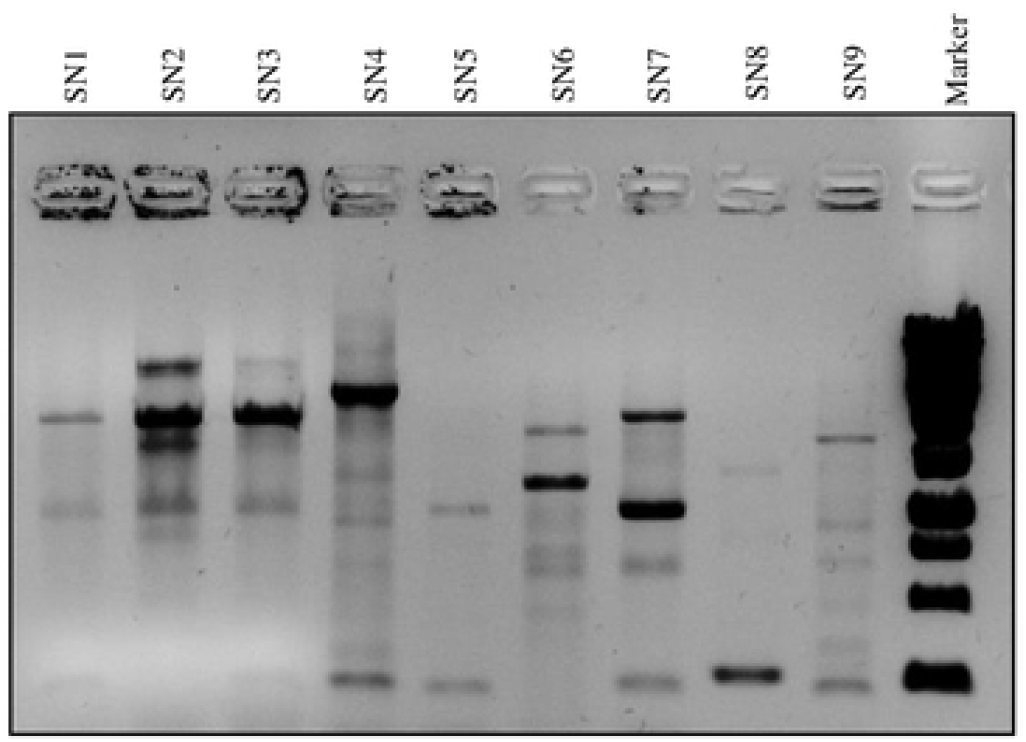
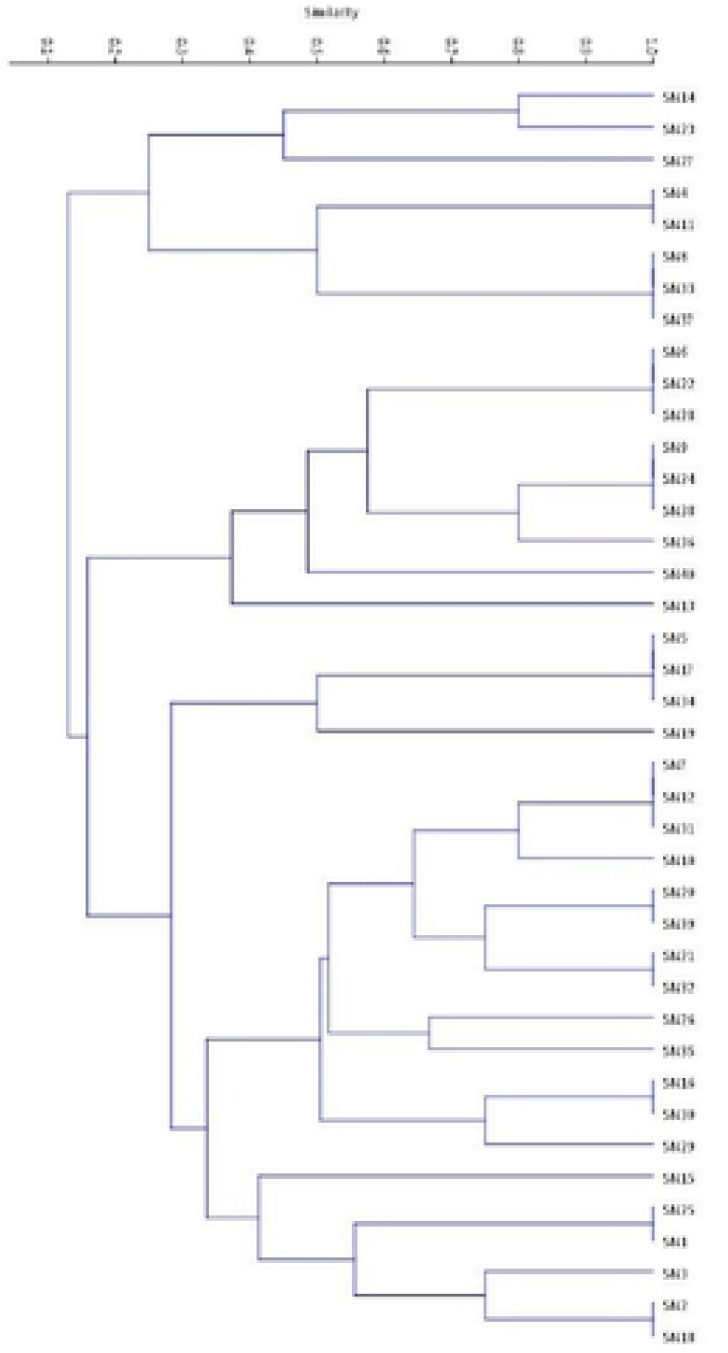
(**a**) **AP-PCR patterns of *S. mutans* isolates**. Colony PCR was performed with the primer set OPA-02 and OPA-13 primers (lanes 1-9). Lane 10 contains the DNA ladder (1 kbp plus). (**b**). **Dendrogram delineating the genetic diversity of the 40 isolated *S. mutans* strains**. The Dice coefficient was computed based on UPGMA clustering algorithm.

### Growth kinetics of *S. mutans* clinical isolates

To investigate the growth kinetics of various clinical isolates, we performed the growth curve analysis for 12 hours in compare to the reference strain UA159. We observed noticeable variation in growth rate in some strains, however, most of the strains grew at similar rate as like UA159 (Fig 2a). Some of the isolates demonstrated very slow growth rate with long dividing time (>150 minutes) and took three to four days to have distinct colony on the agar plate at microaerophilic condition. Similarly, final growth at OD630 after 24-hour incubation was also less for the slow grower (Fig 2b). Growth pattern of the UA159 was in the middle among the isolated strains (dividing time 57 minutes and final growth at OD630 was 1.768).

**Fig 2.**
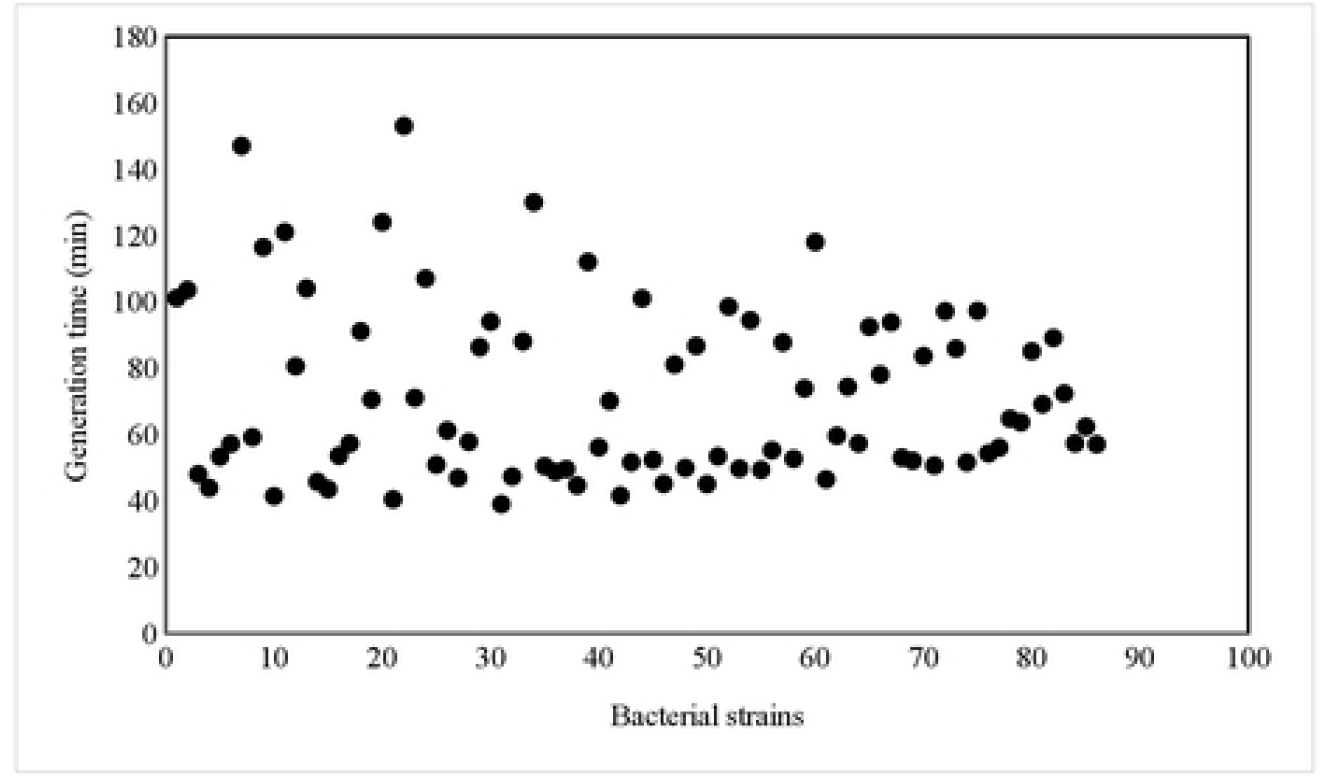
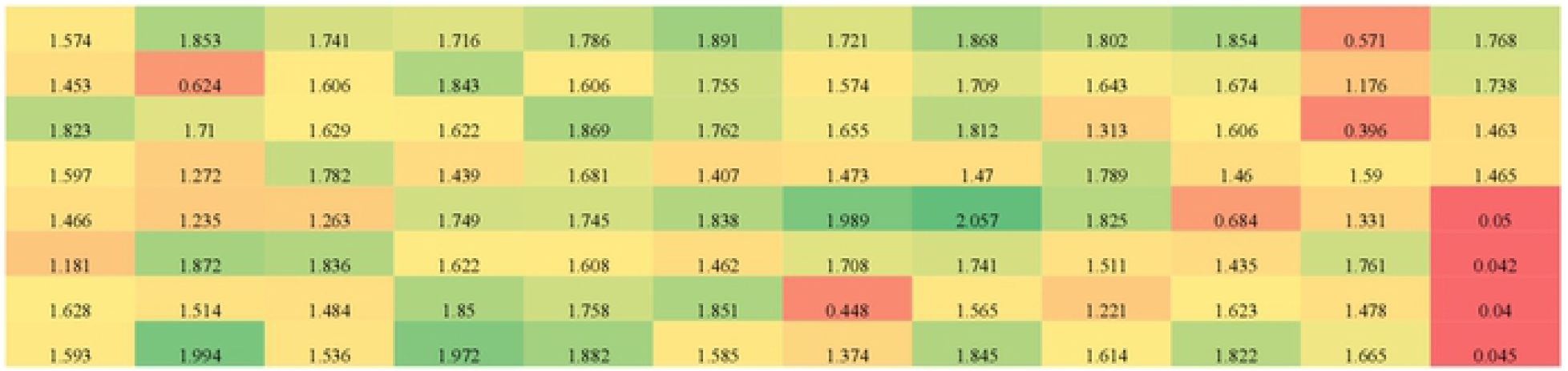
**(a). Mean doubling time of the isolated strains**. Isolates were grown overnight and subcultured to fresh THY broth and absorbance was measured every one-hour interval. Doubling time was calculated based on two OD values taken from the logarithmic phase of the growth by using the formula, r = ln [OD2/OD1]/(T2-T1) and represents the average value of at least two measurements. **(b) Final growth yield of the isolated strains**. Isolates were grown overnight and subcultured to fresh THY broth and absorbance was taken after 24 hours. Each experiment was performed at least twice in duplicates.

### Distribution of *S. mutans* putative virulence genes

We investigated the presence or absence of 15 chromosomally encoded *S. mutans* virulence genes by PCR. Genes involved in various functions such as two component system, bacteriocin production, biofilm formation, acid tolerance, and collagen binding were investigates in this study (Table 3). Eight of these genes were present in all clinical isolates and five genes were differentially present among the isolates. However, collagen binding protein, *cnm* and *cbp*, were not detected in the isolates. The presence of bacteriocin encoding genes (*nlmAB* and *nlmC*) was observed in 63% and 48% of the isolates. We also studied the distribution of two component system ComDE and CiaHR and we found that both systems are present in all of the isolated strains.

**Table 3:**
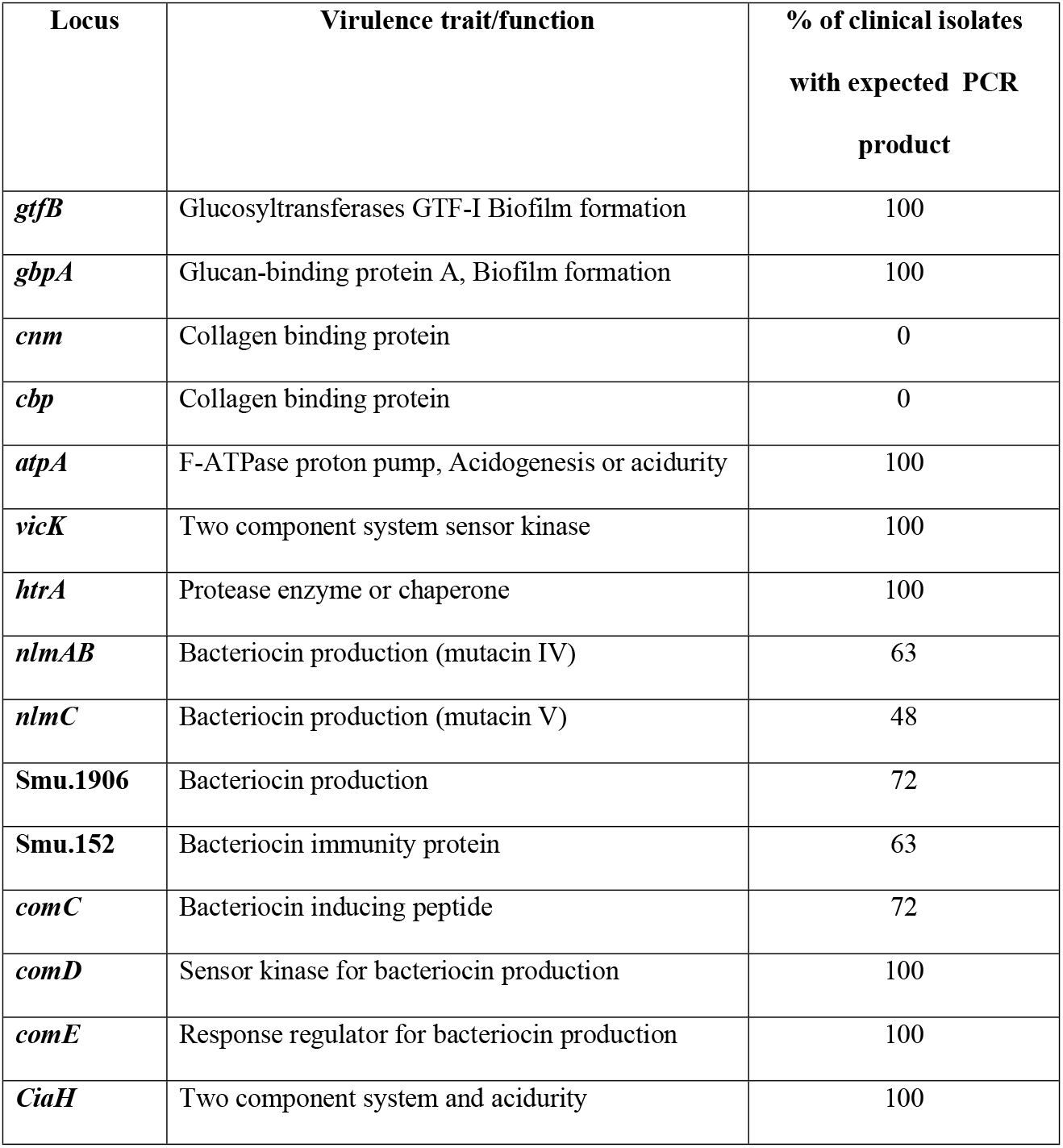
Distribution of virulences genes among *S. mutans* clinical isolates.

### Acid tolerance by *S. mutans* clinical isolates

The ability to grow at low pH is an important virulence attribute for *S. mutans*. To investigate the acid tolerance of S*. mutans* clinical isolates, we cultured the isolated strains in the medium which was acidified at pH 5.0 with HCl. We observed noticeable variation in growth pattern among the isolated strains with the mean generation times ranging from 83 minutes to 234 minutes (Fig 3). Some strains grew faster than UA159.

**Fig 3.**
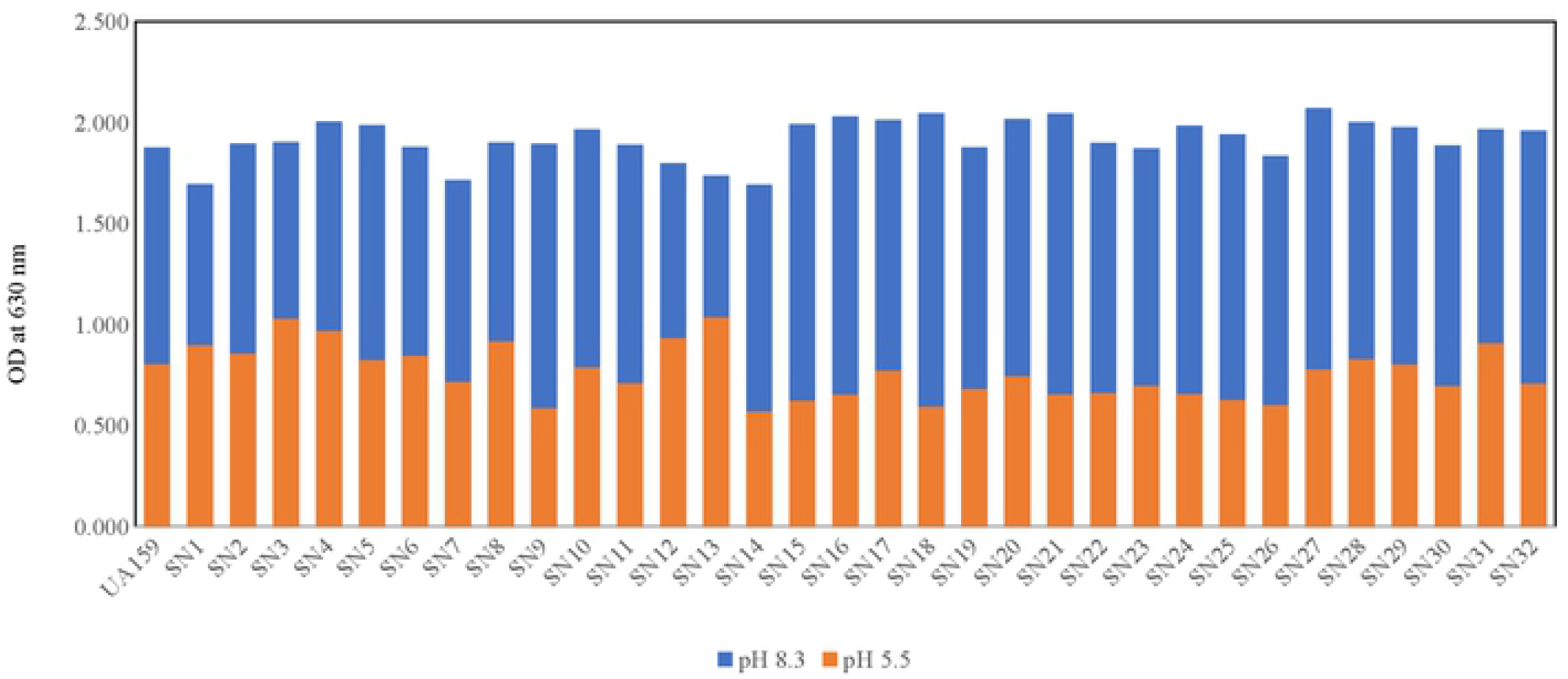
Acid tolerance of the isolated strains. Isolated strains were incubated in THY broth either at pH 8.3 or pH 5.5 and growth was monitored for 24 hour period. Error bar represents the standard deviation.

### Acidogenesis of *S. mutans* clinical isolates

Acidogenesis is the most important virulence factor for dental caries. To investigate the acid production capacity of the clinical isolates, we inoculated the overnight grown *S. mutans* culture in THY broth and measured the pH value at different time intervals. We noticed that all of the clinical isolates have remarkable ability to produce acid while growing in THY broth (Table 2 and Fig 4) and turned the initial pH of the media from 8.34 to more acidic pH (up to pH 5.0). Most of the clinical isolates displayed better acid production ability than the reference strain, UA159 which has turned the THY broth from pH 8.34 to pH 6.25.

**Fig 4.**
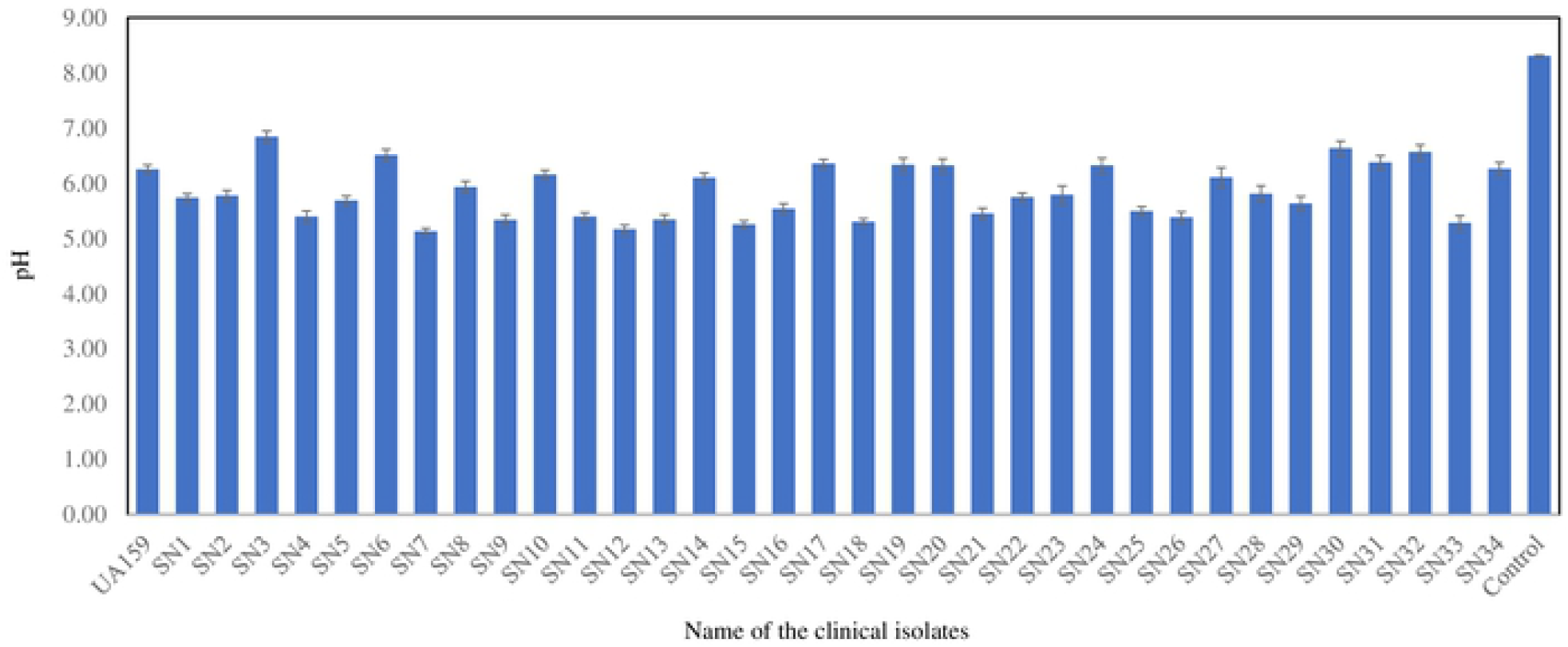
Acid production by the isolated strains. Isolated strains were incubated in THY broth and the pH was measured every 24-hour intervals with a pH meter. Error bar represents the standard deviation.

### Biofilm forming capacity of *S. mutans* clinical isolates

*S. mutans* has the outstanding ability to form biofilm on the teeth surface and causes plaque formation. In order study biofilm forming capacity of various clinical isolates, we cultured them in THY medium supplemented with 1% sucrose for 48 hours in 24 well plates. Our in vitro biofilm formation assay suggested that all the strains retained significant level of biofilm formation capacity as like UA159. However, variation in biofilm formation capacity was also present among the clinical strains and some of the isolates displayed superior biofilm forming capacity than UA159 (Table 2 and Fig 5).

**Fig 5.**
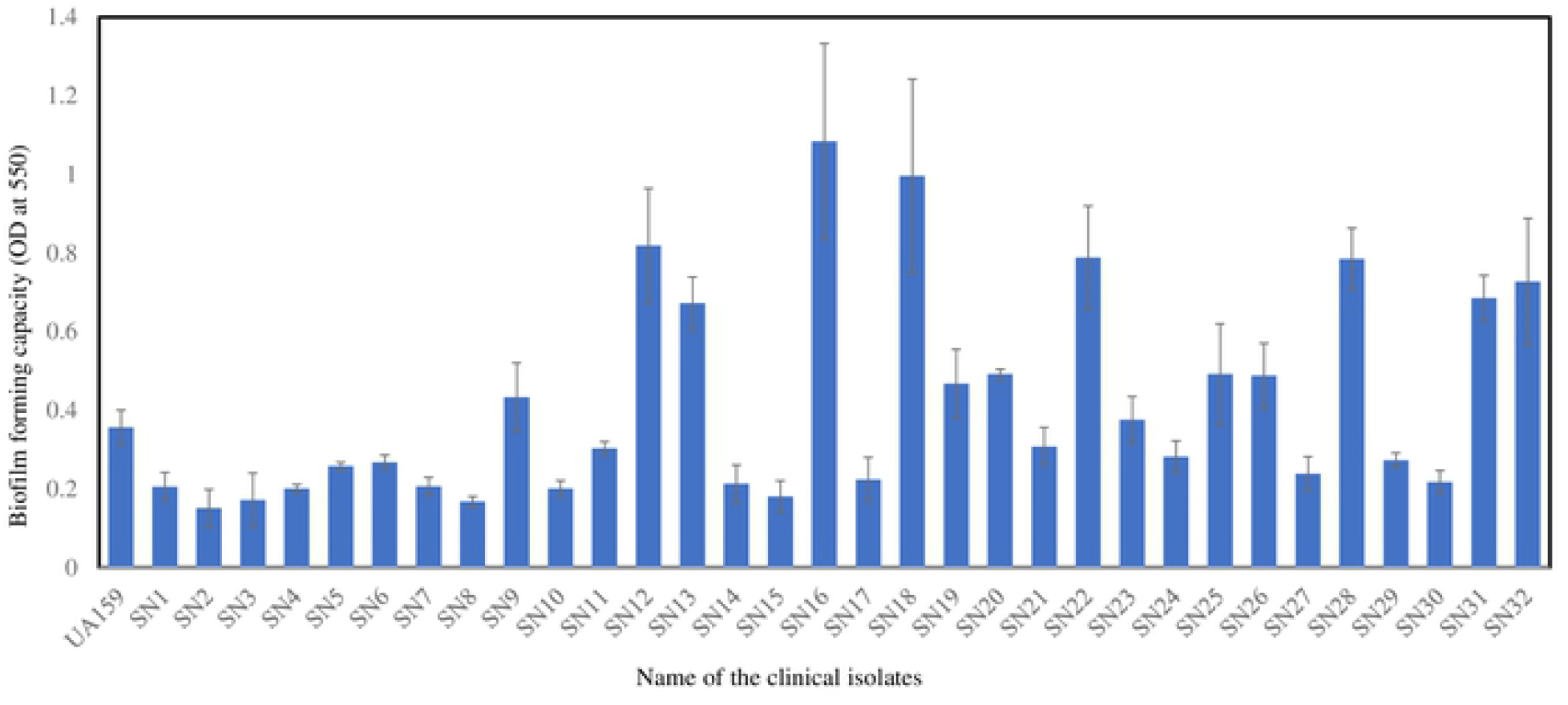
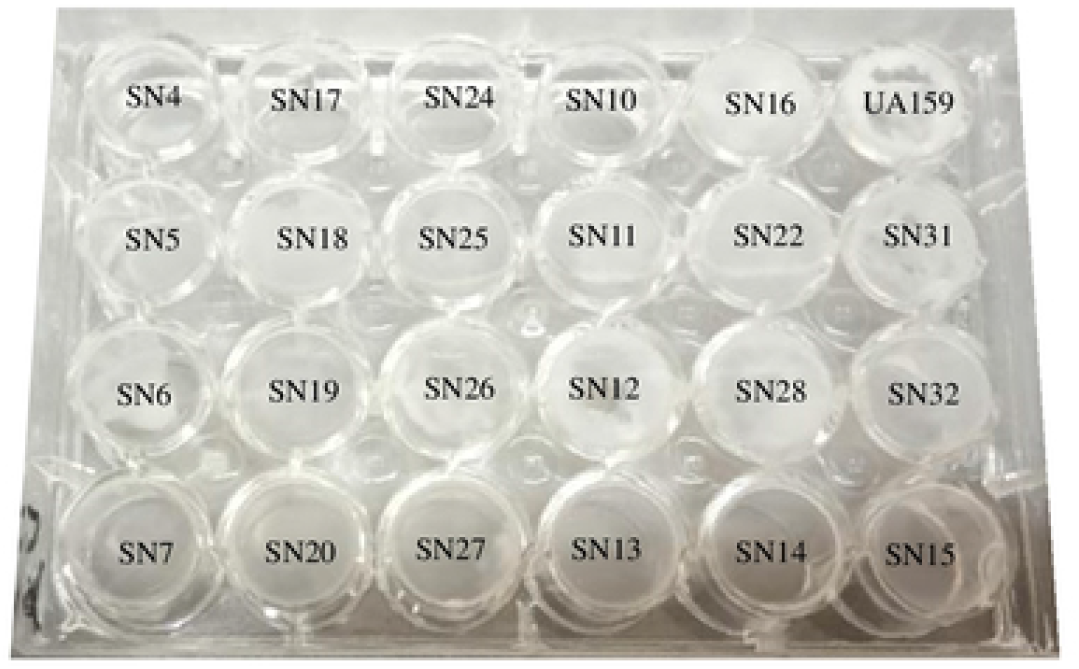
Biofilm formation by the isolated strains. Absorbance values were taken at 550 nm after 48-hour incubation. Error bar represents the standard deviation. Each experiment was performed in triplicates.

### Bacteriocin production by different clinical isolates

*S. mutans* has the capacity to produce various types of bacteriocins to inhibit other competitor microorganisms present in the oral habitat (5). To investigate the ability of different *S. mutans* clinical isolates to produce bacteriocins, we screened all *S. mutans* isolates against *S. pyogenes* (locally isolated) and *L. lactis* (locally isolated). Our study revealed that 17% and 12% of the clinical isolates were able to secrete bacteriocin against *S. pyogenes* and *L. lactis* respectively (Table 2 and Fig 6). Whereas, *S. mutans* UA159 produced antagonistic activity against both indicator bacteria.

**Fig 6.**
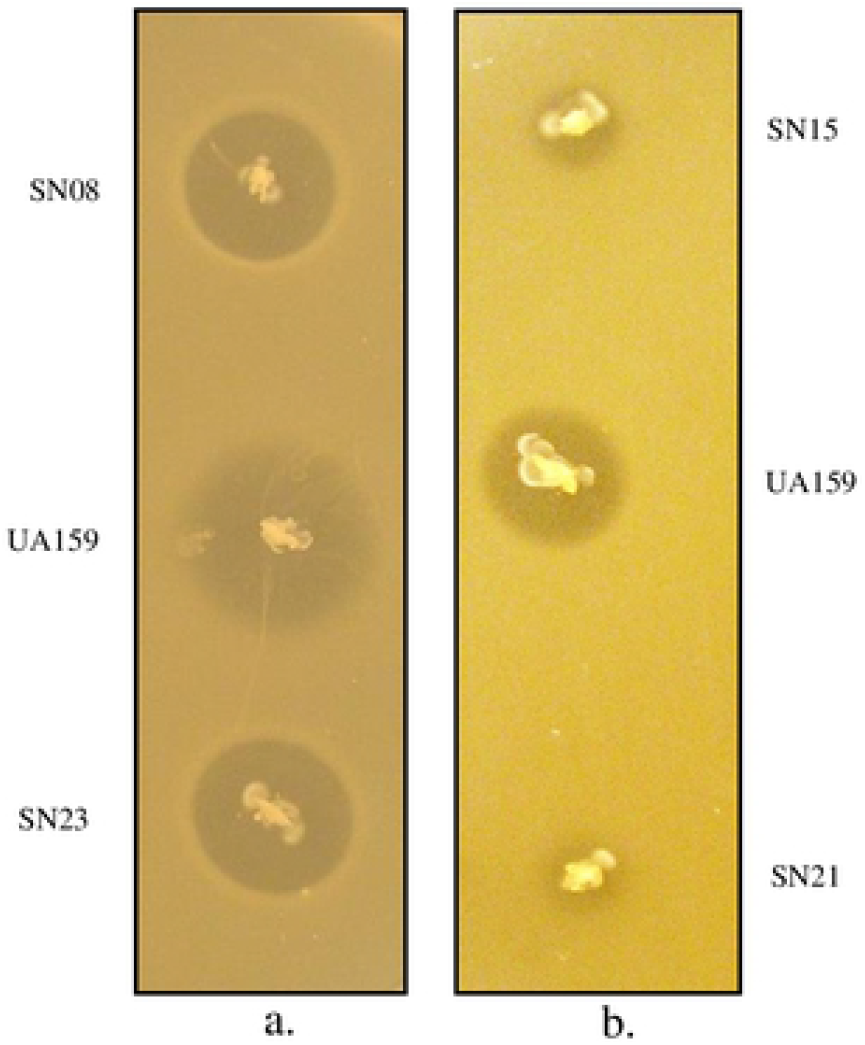
Bacteriocin production by the clinical isolates. A single colony of the clinical isolates were stabbed into the THY agar and incubated for 24-hour. Indicator bacteria were grown overnight and overlaid on the THY agar plate seeded with the clinical isolates as soft agar. Overlaid plates were then incubated overnight under same conditions and the diameter of the zones of inhibition around the producer bacteria was measured. Assays were repeated at least two times and a representative plate is shown.

## Discussion

Oral health is generally considered as a mirror of one’s general health and sometimes associated with several systemic diseases (52). Dental caries, commonly known as tooth decay, is the most common oral health problem worldwide and *S. mutans* is considered to be the primary causative agents of dental caries (53). *S. mutans* resides in the dental plaque, a multispecies biofilm community that harbors more than 700 different types of microorganisms (54). As the biofilm matures, the pioneer colonizers, which are comprised mostly of mitis streptococci, are replaced with early colonizers, such as *S. mutans* (55). The ability to establish biofilm lifestyle, production of organic acid and ability to survive at low pH, outstanding ability to outcompete other bacteria by the production of bacteriocin and generation of genetic diversity by natural transformation are attributed as the prime driving force for its ability to adapt and survive in the rapidly changing environment of the oral cavity (12–16, 19). In this study, we show that the phenotypic and genotypic properties that are associated with the virulence of *S. mutans* are diverse and vary significantly among 209 newly isolated clinical strains.

We found that types and colony morphology of the isolated strains on mitis salivaris agar vary considerably from patient to patient. We observed multiple strains in the same sample with various colony morphology, which further confirmed that dental plaques indeed contain multispecies biofilm structures and intimate association of all the species are required for causing dental caries. We observed *S. mutans* like colony morphology in 82% of the samples and further screening of by species-specific PCR demonstrated that only 76% of the preliminarily selected colonies were *S. mutans*. Among the selected strains, serotype c was found to be the most abundant (77%), followed by serotype e (18%) and serotype f (2%) (data not shown). We did not find any serotype k strains and were unable to serotype 3% of the cases by our molecular approach. Our results are in consistent with other results where serotype c and e shown to be the two abundant serotypes in the dental cavity globally (22, 24). Our results are in consistent with previous results where serotype c was shown to be most prevalent *S. mutans* in the oral cavity (7, 56).

We also performed AP-PCR of 40 selected strains to investigate the genotypic diversity of the isolated strains and found that 26 different genotypes are present among the strains. High levels of genotypic variations were also found previously by several groups (21, 57, 58). Zhou et al. classified 730 *S. mutans* isolates into 337 distinct genotypes by AP-PCR fingerprint analysis (57). In a study with young adults, Emanuelsson et al. (59) noticed only seven genotypes in subjects who had previously experienced dental caries. Napimoga et al. found eight genotypes in caries-active subjects using AP-PCR (60). However, it has been reported that children harbor only one to five distinct genotypes of *S. mutans* (21). The high prevalence of genotypic variations can be attributed to diversified horizontal gene transfer, various nutritional behavior, and chemical environment in the oral cavity.

In accordance with the genotypic diversity, our phenotypic studies revealed that isolated *S. mutans* strains have wide variation in phenotypic diversity. Ability to form biofilm, to sustain the growth at low pH and production of acids are considered as key virulence factors in *S. mutans* and were studied extensively (1). In consistent with previous reports, we also observed high variability of these virulence factors among the isolated strains. Some strains displayed better sucrose-dependent biofilm forming capacity than the universal reference strain, UA159 (Figure 5) and some were crippled in biofilm forming capacity. Biofilm formation capacity of *S. mutans* is aided by various genes, which encode several surface antigens to attach the teeth surface (61, 62). The variation in biofilm forming capacity can be due to the presence or absence of various biofilm associated genes, prevalence of polymorphism among these genes and differential epigenetic regulation. We also investigated the presence of biofilm associated gene, *gbpA* in the clinical isolates, however, this gene was present in all strains. In previous studies, it was shown that strains having recombination in *gtfB* and *gtfC* genes are responsible for poor sucrose-dependent biofilm formation in some strains of *S. mutans* (63, 64). In another study, Nakano et al. also found that *gbpA* gene is absent in some *S. mutans* strains (65). We also investigated the growth kinetics of the isolated strains (Fig 2) and found a wide variation in growth kinetics pattern among the strains. Acid resistance of *S. mutans* strains is conferred by the F_1_F_O_-H^+^-translocating ATPase and the activity and optimum pH of this ATPase enzymes are correlated with acid tolerance of oral bacterium (66). For instance, lactobacilli which are strong aciduric organism exhibit better activity and lower pH optima for the ATPase than the acid-sensitive species, *S. sanguinis* (67). Our results suggest that isolated strains have differential response at acid stress. Most of the clinical strains suffered from growth constraints and individual strain exhibited distinct growth kinetics at pH 5.5 (Fig 3). In addition to acidurity, we also investigated the acidogenesis character of the isolated strains and found noticeable variation among acid production while growing on THY media. However, all the strains could turn the initial medium pH of 8.32 to more acidic pH from 5.02 to 6.54. A large proportion of the clinical isolates displayed better acid production than the UA159 although some of the strains were either equal or poor acidogenic as like UA159. Our results showed little variation from a previous study where equal acidogenecity was observed among the *S. mutans* isolates (56). The apparent variation in acid production may be due to different methods and growth medium used in the studies. Instead of using THY broth at microaerophilic conditions, that study used Phenol Red Dextrose broth (Difco) supplemented with 1 % glucose and anaerobic incubation. When we assessed the correlation of acid production with status of dental caries, we did not find any correlation between acidogenecity and tooth decay status (data not shown). Production of bacteriocins to inhibit closely related bacteria are assumed to be important virulence attributes in *S. mutans*, which encodes several bateriocin encoding genes to inhibit the growth of various bacteria in vitro (13). Mutacin IV and V are two important non-lantibiotic bacteria produced by *S. mutans* to inhibit *S. pyogenes, S. gordonii*, *S. oralis*, *Lactococcus lactis* and other streptococci (5). In this study, we tested the clinical isolates against *S. pyogenes* and *L. lactis* by deferred antagonism bacteriocin assay and observed a wide variation in bacteriocin production. Although a minor fraction (17%, and 12% respectively) was able to display antagonistic activity, both *nlmAB* and *nlmC* genes were present in majority of the isolates (63% and 48% respectively). Previous genetics and biochemical study indicated that the buildup of a processed form of *comC* gene product (CSP) results in activation of a two-component system (ComD and ComE), which induces the expression of bacteriocins (13). In this study, we investigated the presence or absence of *comC, comD*, and *comE* genes among the clinical isolates and found that *comDE*two component systems is present in all the isolates. However, *comC* was absent in 28% of the strains, which might be due to either absence of this gene in the isolates or presence of different version of this gene which was not amplified by the primer sequences. Previously, it has been found that significant numbers of the sequenced strains either lack the *comCDE* genes or contained various mutations that could lead to failure to produce functional ComCDE proteins (68, 69). Polymorphisms within the *comCDE* locus of *S. mutans* isolates have resulted in variation in phenotypic properties associated with deletion of *comDE* in different *S. mutans* strains (69). Our results further confirmed the findings that tremendous variation prevails among *S. mutans* strains in the pathways involved in quorum sensing and bacteriocin production.

In this study, we also investigated the distribution of several putative virulence genes with an aim to identify the genetic elements associated with observed phenotypes. Although the presence or absence of the genetic elements tested did not correlate with caries status, their distribution was strongly associated with the virulent phenotypes. For example, strains lacking *nlmAB, nlmC* or *comC* genes were unable to display antagonistic activity against the indicator bacteria. In an agreement with several previous studies, we also observed significant intraspecies genetic diversity for several genes (12, 17, 52 (Table 3). However, most of the strains are genetically homogenous for genes associated with two-component systems which is in accordance with a previous report which reported a significant level of genetic homogeneity among *S. mutans* strains (29). However, our results are in contrast with Palmer et al. who reported that wide variations exist among strains of *S. mutans* in the pathways involved in quorum sensing, genetic competence and non-lantibiotic bacteriocins (1).

Variation in the same species is prevalent in several bacterial species, either by sharing genes by some but not all isolates or by strain-specific genes that are unique to each isolates (29, 70). In a genome wide comparison, it was revealed that *S. mutans* strains, UA159 and NN2025, differ in 10% of the genes (43) and 20% of the open reading frames (ORFs) in universal reference strain, UA159 have been shown to be dispensable genome by a DNA hybridization-based comparison with nine other strains (71). In addition, another comparative genome hybridization study comprised of 11 strains showed that 16.6% of the ORFs included in the microarray were not present at least one of the genomes (41). Most of the common genes which were present in all the strains in our study were involved in central carbon metabolism and two component systems (Table 3). Our results are in consistent with a previous study where Argimon et al.(29) also observed the widespread present of these genes among 33 *S. mutans* isolates. Intra-species variation in gene content has been reported previously various *S. mutans* genes. For instance, the *gbpA* gene, encoding a glucan-binding protein, was previously found to be absent in five isolates from a collection of 39 laboratory and clinical strains (45). In contrast with this report and in accordance with two previous studies, we found that all the strains carry *gbpA* gene (Table 3) (29, 41). Bacteriocin encoding genes, *nlmAB*, *nlmC*, were present in 63 and 48% of the isolates, respectively in our (Table 3). This is in agreement with previous studies, which found the mutacin IV encoding *nlmA* and *nlmB* genes in 50% of a population of 70 clinical isolates (31) and *nlmC* was found in 60% of the isolates (19, 29). We did not find any *cnm* genes, encodes the collagen-binding protein, among the isolated strains, which is inconsistent with a previous report where *cnm* gene was not detected in a collection of 33 clinical isolates (29). However, two recent studies reported the presence of *cnm* gene in 9.8% (54) and 21.6% (52) of the analyzed strains (40, 44). Taken together all the genotypic and phenotypic assays, our results indicated that *S. mutans* clinical isolates exhibit profound variations among themselves. The distribution of various virulence associated genes is directly correlated with their phenotypes. Further genomics and transcriptomics studies are warranted to get insight into the tremendous variation exists among the *S. mutans* clinical isolates and their possible interactions with symbiotic and antagonistic neighbors prevalent in the oral cavity. It would also be worthwhile to employ metagenomic approaches to understand the complex architecture of dental caries associated microbiome and their ability to cause tooth decay.

The phenotypic and genotypic properties of *S. mutans* clinical isolates presented here imply that clinical strains have undergone intense evolutionary changes to cope up with the rapidly changing environment in the oral cavity. This study has helped us to better understand the cariogenic variations of *S. mutans* clinical strains which can be used to devise new approaches to control *S. mutans* mediated dental diseases thereby. Moreover, the obtained knowledge from this study can be used as a resource to further study the pathogenesis of this bacterium in relation with dental caries and other systemic diseases.

## Acknowledgements

We thank Dr. Shahidul Islam of Bangabandhy Seikh Mujib Medical University for helping us to collect the clinical isolates. This work was supported by research grant provided to Dr. Mohammad Shahnoor Hossain from Biotechnology Research Centre, University of Dhaka Bangladesh.

## Author Contributions

Conceived and designed the experiments: MSH. Analyzed the data: MSH. Wrote the manuscript: MSH. Helps to collect the sample and dental examination: SI. Performed experiments: MSH, SA, YMN, TAT, SMB. Agreed with manuscript results and conclusions: MSH, SA, YMN, TAT, SMB, SI and MSH. Made critical revisions and approved final version: MSH. All authors reviewed and approved the final manuscript.

